# A potential cortical precursor of visual word form recognition in untrained monkeys

**DOI:** 10.1101/739649

**Authors:** Rishi Rajalingham, Kohitij Kar, Sachi Sanghavi, Stanislas Dehaene, James J. DiCarlo

## Abstract

Skilled human readers can readily recognize written letters and letter strings. This domain of visual recognition, known as orthographic processing, is foundational to human reading, but it is unclear how it is supported by neural populations in the human brain. Behavioral research has shown that non-human primates (baboons) can learn to distinguish written English words from pseudo-words (lexical decision), successfully generalize that behavior to novel strings, and exhibit behavioral error patterns that are consistent with humans. Thus, non-human primate models, while not capturing the entirety of human reading abilities, may provide a unique opportunity to investigate the neuronal mechanisms underlying orthographic processing. Here, we investigated the neuronal representation of letters and letter strings in the ventral visual stream of naive macaque monkeys, and asked to what extent these representations could support visual word recognition. We recorded the activity of hundreds of neurons at the top two levels of the ventral visual form processing pathway (V4 and IT) while monkeys passively viewed images of letters, English words, and non-word letter strings. Linear decoders were used to probe whether those neural responses could support a battery of orthographic processing tasks such as invariant letter identification and lexical decision. We found that IT-based decoders achieved baboon-level performance on these tasks, with a pattern of errors highly correlated to the previously reported primate behavior. This capacity to support orthographic processing tasks was also present in the high-layer units of state-of-the-art artificial neural network models of the ventral stream, but not in the low-layer representations of those models. Taken together, these results show that the IT cortex of untrained monkeys contains a reservoir of precursor features from which downstream brain regions could, with some supervised instruction, learn to support the visual recognition of written words. This suggests that the acquisition of reading in humans did not require a full rebuild of visual processing, but rather the recycling of a brain network evolved for other visual functions.

## Introduction

Literate human adults can efficiently recognize written letters and their combinations over a broad range of fonts, scripts and sizes (1–3). This domain of visual recognition, known as orthographic processing, is foundational to human reading abilities, because the invariant recognition of the visual word form is an indispensable step prior to accessing the sounds (phonology) and meanings (semantics) of written words (4). It is largely unknown how orthographic processing is supported by neural populations in the human brain. Given the recency of reading and writing to the human species (a cultural invention dating to within a few thousand years), it is widely believed that the human brain could not have evolved *de novo* neural mechanisms for the visual processing of orthographic stimuli, and that the neural representations that underlie orthographic processing abilities must build upon, and thus be strongly constrained by, the prior evolution of the primate brain (5, 6). In particular, a dominant theory is that the ventral visual pathway, a hierarchy of cortical regions known to support visual object recognition behaviors, could be inherited from recent evolutionary ancestors and minimally repurposed (or “recycled”) through developmental experience to support orthographic processing (6). Consistent with this hypothesis, functional imaging studies suggest that the post-natal acquisition of reading is accompanied by a partial specialization of dedicated cortical sub-regions in the human ventral visual pathway, which ultimately become strongly selective to orthographic stimuli (7–9). However, given the limitations of human imaging methods, it has been challenging to quantitatively test if and how neural representations in the ventral visual pathway might be reused to support orthographic processing.

Interestingly, the ventral visual processing stream – a hierarchically-connected set of neocortical areas (10) – appears remarkably well conserved across many primate species, including Old World monkeys, such as a rhesus macaques (Macaca mulatta) and baboons (Papio papio), that diverged from humans about 25 million years ago (11). Indeed, decades of research have inferred strong anatomical and functional homologies of the ventral visual hierarchy between humans and macaque monkeys (12–14). Previously, we observed striking similarities in invariant visual object recognition behavior between these two primate species, even when measured at very high behavioral resolution (15, 16). Recent work suggests that non-human primates may also mimic some aspects of human orthographic processing behavior (17, 18). In particular, Grainger and colleagues showed that baboons can learn to accurately discriminate visually-presented four-letter English words from pseudo-word strings (17). Crucially, the authors showed that baboons were not simply memorizing every stimulus, but instead had learned to discriminate between these two categories of visual stimuli based on the general statistical properties of English spelling, as they generalized to novel stimuli with above-chance performance. Furthermore, the baboons’ patterns of behavioral performance across non-word stimuli was similar to the corresponding pattern in literate human adults, who make infrequent but systematic errors on this task. Taken together, those prior results suggest that non-human primate models, while not capturing the entirety of human reading abilities, may provide a unique opportunity to investigate the neuronal mechanisms underlying orthographic processing.

In light of this opportunity, we investigated the existence of potential neural precursors of visual word form recognition in the ventral visual pathway of untrained macaque monkeys. Prior neurophysiological and neuropsychological research in macaque monkeys point to a central role of the ventral visual stream in invariant object recognition (19–21), with neurons in inferior temporal (IT) cortex, the top-most stage of the ventral stream hierarchy, exhibiting selectivity for complex visual features and remarkable tolerance to changes in viewing conditions (e.g. position, scale, and pose) (19, 22, 23). It has been suggested that such neural features could have been coopted and selected by human writing systems throughout the world (5, 6, 24). Here, we reasoned that if orthographic processing abilities are supported by “recycling” primate IT cortex – either by minimal adaptations to the IT representation and/or evolutionary addition of new cortical tissue downstream of IT – then this predicts that the initial state of the IT representation, as measured in naïve macaque monkeys, should readily serve as a computational precursor of orthographic processing tests. Investigating, for the first time, the representation of letters and letter strings in macaque IT cortex would not only directly test this prediction but could also provide initial insights into the representation of letters and letter strings prior to reading acquisition.

To quantitatively test this prediction of the “IT precursor” hypothesis, we first operationally defined a set of thirty orthographic identification and categorization tasks, such as identifying the presence of a specific letter or specific bigram within a letter string (invariant letter/bigram identification), or sorting out English words from pseudo-words (lexical decision). We do not claim this to be an exhaustive characterization of orthographic processing, but an unbiased starting point for that greater goal. As schematically illustrated in Figure 1A, we then recorded the spiking activity of hundreds of neural sites in V4 and IT of rhesus macaque monkeys while they passively viewed images of letters, English words and non-word strings. We then formally tested this prediction of the IT precursor hypothesis by asking whether adding a simple neural readout machinery on top of the macaque IT representation could produce a neural substrate of orthographic processing, using biologically plausible linear decoders that perform those behavioral tasks from the firing responses of those neuronal populations. We found that linear decoders that learn from the population spiking output of IT cortex easily achieved baboon-level performance on these tasks, and that the pattern of behavioral performance predicted by this hypothesis was highly correlated with the corresponding baboon behavioral pattern. These behavioral tests were also met by leading artificial neural network models of the non-human primate ventral stream, but not by low-level representations of those models. Taken together, these results show that, even in untrained non-human primates, the population of IT neurons contains an explicit (i.e. linearly separable), if still imperfect, representation of written words that might have been later “recycled” to support orthographic processing behaviors in higher primates such as humans.

**Figure 1:**
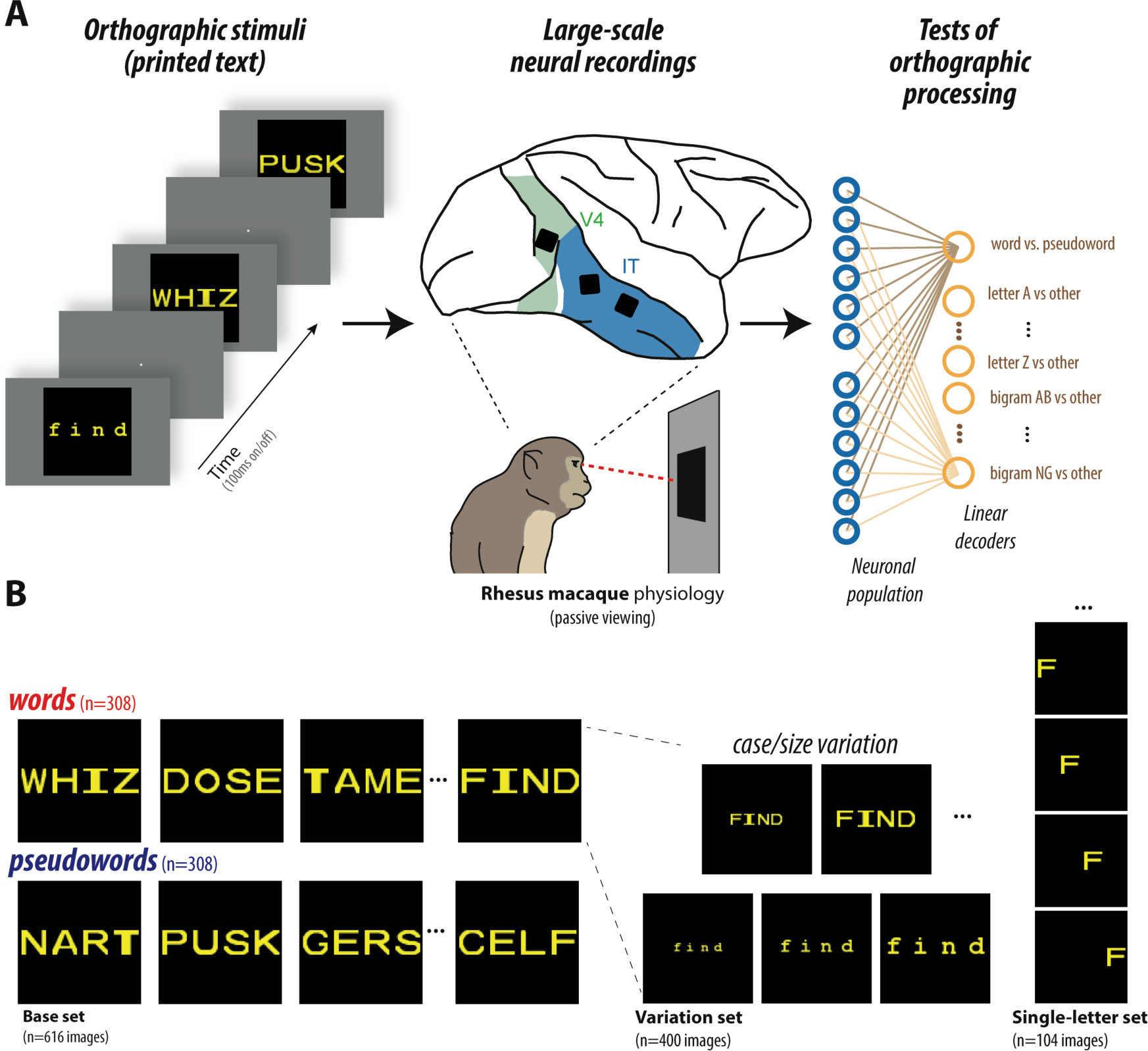
**(A)** Schematic of experiment. We recorded the activity of hundreds of neural sites in IT while monkeys passively fixated images of orthographic stimuli. (As a control, we also recorded from the dominant input to IT, area V4.) We then tested the sufficiency of the IT representation on 30 tests of orthographic processing (e.g. lexical decision, letter identification, etc.) using simple linear decoders. **(B)** Example visual stimuli. Images consisted of four-letter English words and pseudo-word strings presented in canonical views, as well as with variation in case (upper/lower) and size (small/medium/large), and single alphabet letters presented at four different locations.

## Results

Our primary goal was to experimentally test the capacity of neural representations in the primate ventral visual pathway to support orthographic classification tasks. To do so, we recorded the activity of hundreds of neurons from the top two levels of the ventral visual cortical hierarchy of rhesus macaque monkeys. Neurophysiological recordings were made in four Rhesus monkeys using chronically implanted intracortical microelectrode arrays (Utah) implanted in the inferior temporal (IT) cortex, the top-most stage of the macaque ventral visual stream hierarchy (IT). As a control, we also collected data from upstream visual cortical area V4, which provides the dominant input to IT (Figure 1A). Neuronal responses were measured while each monkey passively viewed streams of images, consisting of alphabet letters, English words, and pseudo-word strings, presented in a rapid serial visual presentation (RSVP) protocol at the center of gaze (Fig. 1). Images were presented in randomized order, and each image was shown at least 25 times. Crucially, monkeys had no previous supervised experience with orthographic stimuli, and they were not rewarded contingently on the stimuli, but solely for accurately fixating. This experimental procedure resulted in a large dataset of 510 IT neural sites (and 277 V4 neural sites) in response to up to 1120 images of orthographic stimuli. To test the sufficiency of the IT representation for orthographic processing, we used simple linear decoders (as biologically plausible approximations of downstream neural computations, see Methods) to test each neuronal population on a battery of 30 visual orthographic processing tasks: 20 invariant letter identification tasks, 8 invariant bigram identification tasks, and two variants of the lexical decision task. For each behavioral test, we used a linear decoder, which computes a simple weighted sum over the IT population activity, to discriminate between two classes of stimuli (e.g. words versus pseudo-words). The decoder weights are “learned” using the IT population responses to a subset of stimuli (using 90% of the stimuli for training), and then the performance of the decoder is tested on held-out stimuli. The overarching prediction of the “IT precursor” hypothesis was that, if a putative neural mechanism (i.e. a particular readout of a particular neural population) is sufficient for primate orthographic processing behaviors, then, it should be easy to learn (i.e. few supervised examples), its learned performance should match the overall primate performance, and its learned performance should have similar patterns of errors as primates that have learned those same tasks. This logic has been previously applied to the domain of core object recognition to uncover specific neural linking hypotheses (25) that have been successfully validated with direct causal perturbation of neural activity (26, 27).

### Lexical decision

We first focused on the visual discrimination of English words from pseudo-words (a.k.a. lexical decision) using a random subset of the stimuli tested on baboons (17). We collected the response of 510 IT neural sites and 277 V4 neural sites to a base set of 308 four-letter written words and 308 four-letter pseudo-words (see Figure 1B, “base set” for example stimuli). To test the capacity of the IT neural representation to support lexical decision, we trained a linear decoder using the IT population responses to a subset of words and pseudo-words, and tested the performance of the decoder on held-out stimuli. Note that this task requires generalization of a learned lexical classification to novel orthographic stimuli, rather than the mere memorization of orthographic properties. Figure 2A shows the output choices of the linear readout of IT neurons, plotted as the probability of categorizing stimuli as words, as compared to behavioral choices of a pool of six baboons, as previously measured by Grainger et al. (17). For ease of visualization, the 616 individual stimuli were grouped into equally-sized bins based on the baboon performance, separately for words and pseudo-words. We qualitatively observe a tight correspondence between the behavioral choices made by baboons and those measured by the linear decoder trained on the IT population. To quantify this similarity, we benchmarked both the overall performance (accuracy) and the consistency of pattern of errors of the IT population with respect to this previously measured median baboon behavior on the same images.

**Figure 2:**
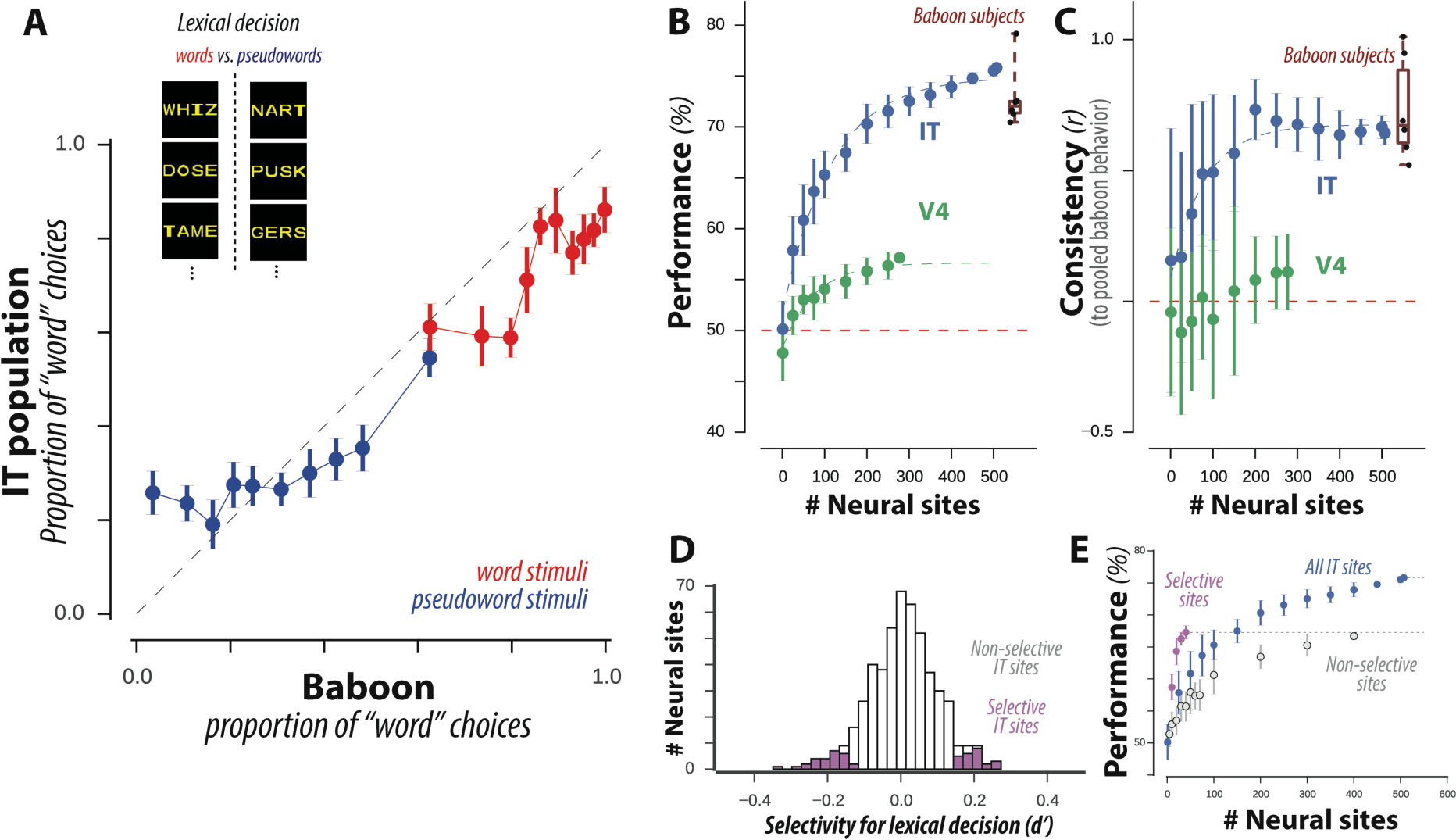
**(A)** Comparison of baboon behavior and a linear readout of IT neurons, plotted as the proportion of stimuli categorized as “words.” The 616 individual stimuli were grouped into equally-sized bins based on the baboon performance, separately for words (red) and pseudo-words (blue). Error bars correspond to SEM, obtained via bootstrap resampling over stimuli; dashed line corresponds to unity line, demarking a perfect match between baboon behavior and IT-based decoder outputs. **(B)** Performance of decoders trained on IT and V4 representations on lexical decision, for varying number of neural sites. Distribution of individual baboon performance is shown on the right. **(C)** Consistency with baboon behavioral patterns of decoders trained on IT and V4 representations, for varying number of neural sites. **(D)** Distribution of selectivity of lexical decision for individual IT sites, highlighting the subpopulation of sites with selectivity significantly different from zero. **(E)** Performance of decoders trained on subpopulation of selective sites from (d) compared to remaining IT sites and all IT sites.

We first found that decoders trained on the IT population responses achieved high performance (76% for 510 neural sites) on lexical decision on new images (Figure 2B). Performance increased steadily with the number of neural sites included in the decoder, with about 250 randomly sampled IT neural sites matching the median performance of baboons doing this task (Figure 2B, blue). Could any neural population achieve this performance? As a first control for this, we tested the upstream cortical area V4. We found that the tested sample of V4 neurons did not achieve high performance (only 57% for 277 V4 neural sites), failing to match baboon performance on this task (Figure 2B, green). Going beyond the summary statistic of average performance, we next tested whether baboons and neural populations exhibited similar *behavioral patterns* across stimuli, e.g. whether letter strings that were difficult to categorize for baboons were similarly difficult for these neural populations. To reliably measure behavioral patterns in each individual baboon subject, we grouped the 616 individual stimuli into equally-sized bins based on an independent criterion (the average bigram-frequency of each string in English, see Methods), separately for words and pseudo-words. For both baboons and decoders, we then estimated the average unbiased performance for each stimulus bin using a sensitivity index (d’); this resulted in a ten-dimensional pattern of unbiased performances. We then measured the similarity between patterns of unbiased performances from a tested neural population and the pool of baboons using a noise-adjusted correlation (see Methods). We observed that the pattern of performances obtained from the IT population was highly correlated with the corresponding baboon pool behavioral pattern (noise-adjusted correlation 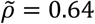; Figure 2C, blue). Perhaps any neural population would exhibit this baboon-like behavioral pattern? On the contrary, we found that this correlation was significantly higher than the corresponding value estimated from the V4 population, which is only one visual processing layer away from IT (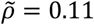; Figure 2C, green). By holding out data from each baboon subject from the pool, we additionally estimated the consistency between each individual baboon subject to the remaining pool of baboons (median 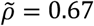, inter-quartile range = 0.27, n=6 baboon subjects). This consistency value corresponds to an estimate of the ceiling of behavioral consistency, accounting for inter-subject variability. Importantly, the consistency of IT-based decoders is within this baboon behavioral range; this demonstrates that that this neural mechanism is as consistent to the baboon pool as each individual baboon is to the baboon pool, at this behavioral resolution. Together, these results suggest that the distributed neural representation in macaque IT cortex is sufficient to explain the lexical decision behavior of baboons, which itself was previously found to be correlated with human behavior (17).

We next explored how the distributed IT population’s capacity for supporting lexical decision arose from single neural sites. Figure 2D shows the distribution of word selectivity of individual sites in units of d’, with positive values corresponding to increased firing rate response for words over pseudo-words. We observed that, across the population, IT did not show strong selectivity for words over pseudo-words (average d’ = 0.008 ± 0.09, mean, SD over 510 IT sites), and that no individual IT site was strongly selective for words vs. pseudo-words (|d’|<0.5 for all recorded sites). However, a small but significant proportion of sites (10%; p<10-5, binomial test with 5% probability of success) exhibited a weak but significant selectivity for this contrast (inferred by a two-tailed exact test with bootstrap resampling over stimuli). Note that this subset of neural sites includes both sites that responded preferentially to words and sites that responded preferentially to pseudo-words. We measured the lexical decision performance of decoders trained on this subpopulation of neural sites, compared to the remaining subpopulation. Importantly, to avoid a selection bias in this procedure, we selected and tested neural sites based on independent sets of data (disjoint split-halves over trial repetitions). As shown in Figure 2E, we observed that decoders trained on this subset of selective neural sites performed better than a corresponding sample from the remaining non-selective population, but not as well as decoders trained on the entire population, suggesting that the population’s capacity for supporting lexical decision relies heavily but not exclusively on this small subset of selective neural sites. We next examined whether this subset of selective neural sites was topographically organized on the cortical tissue. For this subset of neural sites, we did not observe a significant hemispheric bias (p=0.13, binomial test with probability of success matching our hemisphere sampling bias), or significant spatial clustering within each 10×10 electrode array (Moran’s I=0.11, p=0.70, see Methods). Thus, we observed no direct evidence for topographically organized specialization (e.g. orthographic category-selective domains) in untrained non-human primates, at the resolution of single neural sites. Taken together, these results suggest that lexical decision behavior could be supported by a sparse, distributed read-out of the IT representation in untrained monkeys, and provide a baseline against which to compare future studies of trained monkeys.

### Tests of invariant orthographic processing

Importantly, human readers can not only discriminate between different orthographic objects, but also do so with remarkable tolerance to variability in printed text. For example, readers can effortlessly recognize letters and words varying in up to two orders of magnitude in size, and are remarkably tolerant to variations in printed font (e.g. upper vs lower case) (3, 28). To investigate such invariant orthographic processing behaviors, we measured IT decoder performance for stimuli that vary in font size and font case, for a subsampled set of strings (40 words, 40 pseudo-words, under five different variations for a total of 400 stimuli). To test this, we trained linear decoders on subsets of stimuli across all variations, and tested on held-out stimuli, for a total of 29 behavioral tests (20 invariant letter recognition tests, 8 invariant bigram recognition tests, and one test of invariant lexical decision). Figure 3A shows the performance of a decoder trained on the IT neuronal representation on each of these three types of behavioral tests, as a function of the neural sample size. For comparison, we also show the same decoder test for the V4 population. Once again, we observe that the IT population achieved relatively high performance across all tasks, and that this performance was greater than the corresponding estimated performance from the measured V4 population. We note that performance values for invariant lexical decision should not be directly compared with those in Figure 2B, as invariant tests here were conducted with fewer training examples for the decoders (i.e. trained/tested on 40 distinct words/pseudo-words strings, rather than 308 strings in Figure 2B).

**Figure 3:**
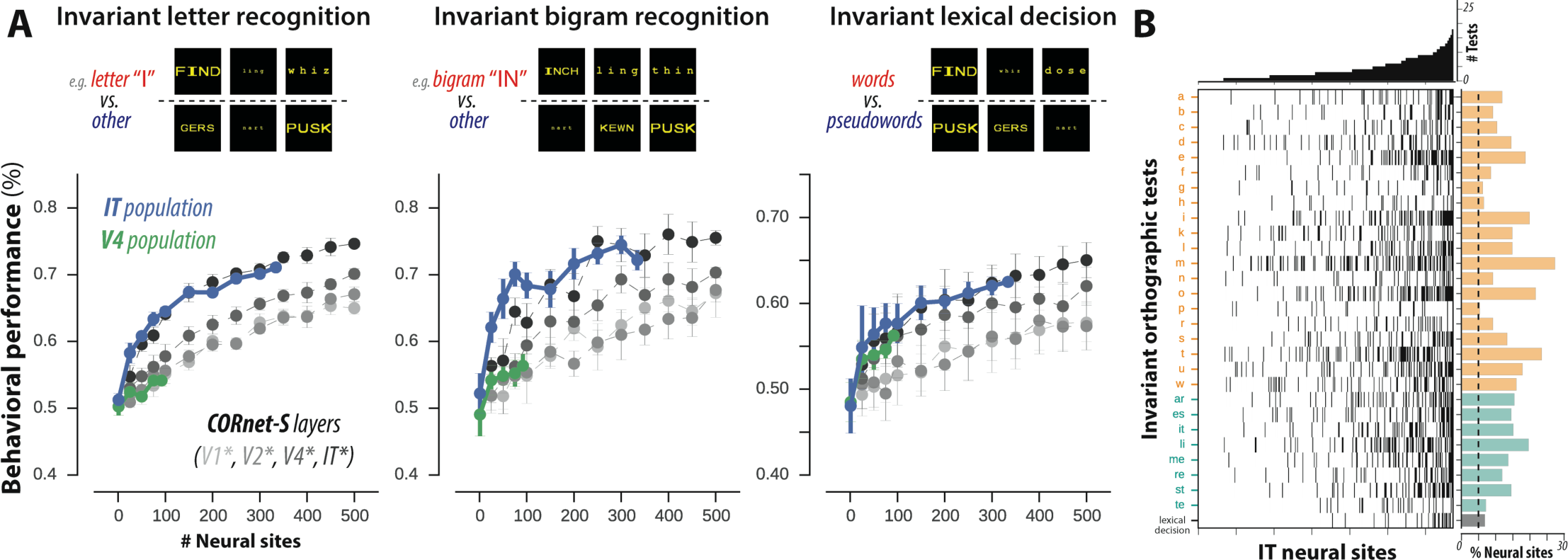
**(A)** Performance of decoders trained on the IT and V4 representations on invariant orthographic tests, grouped into letter identification (n=20 tests), bigram identification (n=8 tests) and invariant lexical decision. Performance of artificial representations sampled from layers of deep convolutional neural network model CORnet-S are shown in grey. **(B)** Selectivity of individual IT sites over 29 invariant orthographic processing tests. The heatmap shows selectivity significantly different from zero over all pairs of neural sites and tests. The histogram above shows the number of behavioral tests (N_t_) that each neural site exhibited selectivity for; neural sites are ordered by increasing N_t_. The histogram on the right shows the proportion of neural sites exhibiting selectivity for each test; the behavioral tests are ordered alphabetically within each task group (letter identification in orange, bigram identification in cyan, and lexical decision in gray). Dashed line corresponds to proportion of tests expected from chance (alpha = 5%).

We additionally tested the feature representation obtained from a deep recurrent convolutional neural network model of the ventral stream on the exact same behavioral tasks. Specifically, we tested the CORnet-S model (29), as it has recently been shown to best match the computations of the primate ventral visual stream (30, 31) and provides an independently simulated estimate of the neuronal population responses from each retinotopically-defined cortical area in the ventral visual hierarchy (V1, V2, V4, and IT). Figure 3 shows the performance of decoders trained on each simulated neuronal population (gray lines) on invariant letter identification, invariant bigram identification, and invariant lexical decision, as a function of the number of model units used for decoding. We observe that the last layer of CORnet-S (simulated IT) significantly outperforms earlier layers (simulated upstream areas V1, V2, and V4) on these invariant orthographic discrimination tasks, and tightly matches the performance of the actual IT population.

Finally, we tested how the IT population’s capacity for these 29 invariant orthographic processing tests was distributed across individual IT neural sites. We computed the selectivity of individual sites in units of d’ for each of these tests, and estimated the statistical significance of each selectivity index using a two-tailed exact test with bootstrap resampling over stimuli (see Methods). Figure 3B shows a heatmap of significant selectivity indices for all pairs of neural sites and behavioral tests; each row corresponds to one behavioral test, each column to a single IT neural site, and filled bins indicate statistically significant selectivity. The histogram above shows the number of behavioral tests that each neural site exhibited selectivity for (median: 3 tests, inter-quartile-range: 5), and the histogram on the right shows the proportion of neural sites exhibiting selectivity for each test (median: 49/337 neural sites, inter-quartile range: 23/337).

Taken together, these results suggest that a sparse, distributed read-out of the adult IT representation of untrained non-human primates is sufficient to support many visual discrimination tasks, including ones in the domain of orthographic processing, and that that neural mechanism could be learned with a small number of training examples (median: 48 stimuli; inter-quartile range: 59, n = 30 behavioral tests). Furthermore, this capacity is not captured by lower-level representations, including neural samples from the dominant visual input to IT (area V4) and low-level ventral stream representations as approximated by state-of-the-art artificial neural network models of the ventral stream.

### Encoding of orthographic stimuli

Finally, the availability for the first time of IT neuronal responses to orthographic stimuli allowed us to begin to address the question of how such stimuli are encoded at the single-neuron level. Behavioral and brain-imaging observations in human readers have led to several proposals concerning the putative neural mechanisms underlying human orthographic abilities. A presumed front end, common to many models, is a bank of letter detectors (e.g.(2, 32, 33)), i.e. a spatially organized array of input units each sensitive to the presence of a specific letter at a given location. Additionally, it has been proposed that written words could be encoded by a set of bigram-sensitive units responding to specific ordered pairs of letters (32). The local combination detector (LCD) hypothesis posits a hierarchy of cortical representations whereby neurons encode printed words at increasing scale and complexity, from tuning to simple edges and letters to intermediate combinations of letters (e.g. letter bigrams) and finally to complex words and morphemes over the cortical hierarchy (2). Other theories have proposed that letter position information is encoded in the precise timing of spikes (34, 35). To date, it has been difficult to directly test such hypotheses. Here, to help constrain the space of encoding hypotheses, we characterized the response properties of hundreds of individual IT neural sites to words and to their component letters.

We first asked if individual IT neural sites exhibit any selectivity for letters. To test this, we measured the selectivity of IT responses to each of the 26 alphabet letters, each presented at four different retinal positions. Figure 4A shows the “tuning curve” for three example IT neural sites. Consistent with the known image selectivity and position tolerance of IT neurons (19, 22, 23), we observed that the responses of these IT neural sites were significantly modulated by both letter identity and letter position, with each example site responding to some but not all individual letters. We focused on 222 (out of 338) neural sites with reliable response patterns across the single letter stimulus set (p<0.01, significant Pearson correlation across split-halves over repetitions). The top panel of Figure 4B shows the average normalized response to each of the 26 letters, across these 222 neural sites. For each neural site, letters were sorted according to the site’s response magnitude, estimated using half of the data (split-half of stimulus repetitions) to ensure statistical independence; we then plotted the sorted letter response measured on the remaining half (individual sites in grey, mean ± SEM in black). Across the entire population, we observe that some neural sites reliably respond more to some letters than others, but this modulation is generally not selective for one or a small number of letters. Rather, sites tended to respond to a broad range of letters, as quantified by the sparsity of letter responses (Figure 4B, bottom panel). Individual sites were also modulated by letter position (Figure 4C, formatted as in Figure 4B), with a greater response to letters presented contralateral to the recording site, while also exhibiting substantial tolerance across positions.

**Figure 4:**
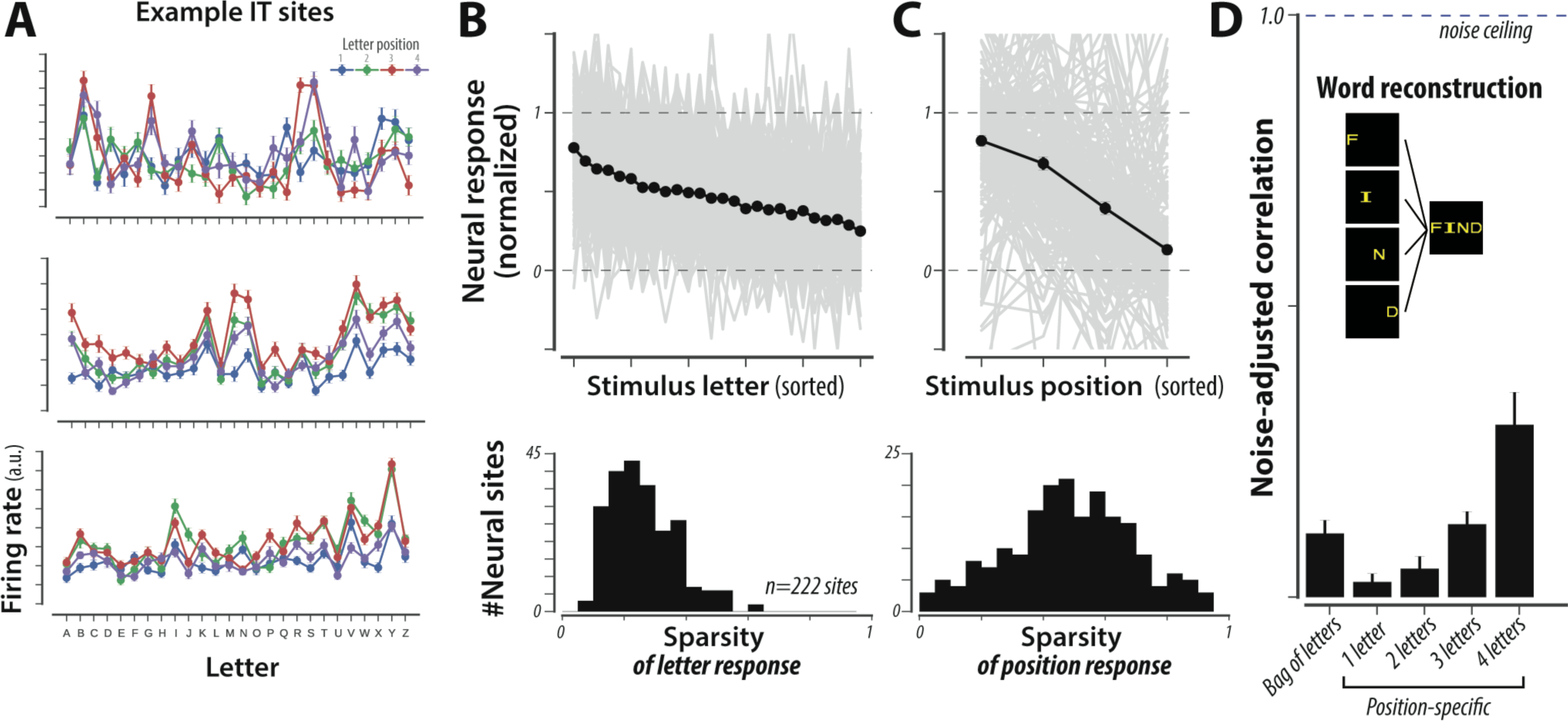
**(A)** Firing rate responses to individual letter stimuli (26 letters at four positions) for three example neurons. **(B)** (top) Average normalized response to each of the 26 letters, across 222 IT neural sites. For each neural site, letters were sorted according to the site’s response magnitude (estimated using an independent half of the data) and plotted in gray. Averaging across the entire population, we observe that neural sites reliably respond more to some letters than others (black, mean ± SE across sites; note that SE is very small). However, this modulation is not very selective to individual letters or small numbers of letters, as quantified by the sparsity of letter responses (bottom panel). **(C)** Individual sites were also modulated by the letter position, exhibiting substantial tolerance across positions (formatted as in B). **(D)** To test if the encoding of letter strings can be approximated as a local combination of responses to individual letters, we reconstructed letter string responses from letter responses, for each neural site. As illustrated by the inset, we used the neural response to images of individual constituent letters to predict the response to images of the corresponding letter strings; predictions were made using linear regressions, crossvalidating over letter strings. The bar plot shows the noise-adjusted correlation of different regression models (median ±SE across neural sites). The “bag of letters” model uses responses of each of the four letters, at arbitrary positions, to predict responses of whole letter strings. Each of the position-specific models uses the responses of up to four letters at the appropriate position to predict letter string responses.

Next, we asked whether the encoding of letter strings could be approximated as a local combination of responses to individual letters. To test this, we linearly regressed each site’s response to letter strings on the responses to the corresponding individual letters at the corresponding position, cross-validating over letter strings. Using the neural responses to all four letters, we observed that the predicted responses of such a linear reconstruction were modestly correlated with the measured responses to letter strings (see Figure 4D, right-most bar; 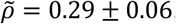, median ± standard error of median, n = 222 neural sites). To investigate if this explanatory power arose from all four letters, or whether 4-letter string responses could be explained just as well by a substring of letters, we trained and tested linear regressions using responses to only some (1, 2, or 3) letters. Given that there are many possible combinations for each, we selected the best such mapping from the training data, ensuring that selection and testing were statistically independent. We observed that reconstructions using only some of the letters were significantly poorer in predicting letter string responses (three letters: 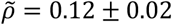, median ± standard error of median). Finally, we tested how well a position-agnostic (or “bag of letters”) model performed on the same reconstruction task by trained and test linear regressions that mapped responses of letters, with the incorrect position (using a fixed, random shuffling of letter positions) on reconstructing the responses to whole letter strings. We found that this “bag of letters” model performed significantly worse (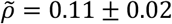, median ± standard error of median).

Note that all correlation values reported above were adjusted to account for the reliability of measured neural responses, such that a fully predictive model would have a noise-adjusted correlation of 1.0 regardless of the finite amount of data that were collected. Yet, the maximal value of 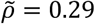 that we obtained using the linear superposition of position-specific responses to the four letters was substantially lower than 1.0. Thus, the pure summation of neural responses to individual letter identity and position explained only a small part of the reliable neural responses to 4-letter strings, suggesting that non-linear responses to local combinations of letters were also present. Future work using stimuli comprising a larger number of letter combinations can explore to what extent IT neural sites respond, for instance, to specific letter bigrams, as predicted by some models (2, 32).

Taken together, these observations demonstrate that individual IT neural sites in untrained non-human primates while failing to exhibit strong orthographic specialization, collectively suffice to support a battery of orthographic tasks. Importantly, these observations establish a number of relevant quantitative baselines, a pre-registered benchmark to which future studies of the ventral stream representations in monkeys trained on orthographic discriminations, or in literate humans, could be directly compared to.

## Discussion

A key goal of human cognitive neuroscience is to understand how the human brain supports the ability to learn to recognize written letters and words. This question has been investigated for several decades using human neuroimaging techniques, yielding putative brain regions that may uniquely underlie orthographic abilities (7–9). In the work presented here, we sought to investigate this behavioral domain in the primate ventral visual stream of naïve rhesus macaque monkeys. Non-human primates such as rhesus macaque monkeys have been essential to study the neuronal mechanisms underlying human visual processing, especially in the domain of object recognition where monkeys and humans exhibit remarkably similar behavior and underlying brain mechanisms, both neuroanatomical and functional (13–16, 36, 37). Given this strong homology, and the relative recency of reading abilities in the human species, we reasoned that the high-level visual representations in the primate ventral visual stream could serve as a precursor that is recycled by developmental experience for human orthographic processing abilities. In other words, we hypothesized that the neural representations that directly underlie human orthographic processing abilities must be strongly constrained by the prior evolution of the primate visual cortex, such that representations present in naïve, illiterate, non-human primates could be minimally adapted to support orthographic processing. Here, we observed that orthographic information was explicitly encoded in sampled populations of spatially distributed IT neurons in naïve, illiterate, non-human primates. Our results are consistent with the hypothesis that the population of IT neurons in each subject forms an explicit representation of orthographic objects, and could serve as a common substrate for learning many visual discrimination tasks, including ones in the domain of orthographic processing.

We tested a battery of 30 orthographic tests, spanning a lexical decision task (words versus pseudo-words), invariant letter recognition, and invariant bigram recognition, as well as modifications that required tolerance to variation in text size and case. We do not claim that these tasks form an exhaustive characterization of orthographic processing, but rather a good starting point for that greater goal. Importantly, this battery of tasks could not be accurately performed by linear readout of the predominant input visual representation to IT (area V4) or by approximations of lower levels of the ventral visual stream, unlike many other coarse discrimination tasks (e.g. contrasting orthographic and non-orthographic stimuli). We tested arbitrarily sampled IT neural sites, including all sampled neural sites with significant visual drive. Finally, we modelled plastic changes via a linear classifier, a simple biologically plausible model of downstream neuronal computations. Indeed, the trained linear decoder performed binary classifications by computing weighted sums of IT responses followed by a decision boundary, analogous to synaptic strengths and spiking thresholds of neurons downstream of IT. We note that the successful classifications we observed do not necessarily imply that the brain exclusively uses IT or the same coding schemes and algorithms that we have used for decoding. Rather, the existence of this sufficient code in untrained and illiterate non-human primates suggests that the primate ventral visual stream could be minimally adapted through experience-dependent plasticity to support orthographic processing behaviors.

These results are consistent with a variant of the “neuronal recycling” theory, which posits that the features that support visual object recognition may have been coopted for written word recognition (5, 6, 24). Specifically, this variant of the theory is that humans have inherited a pre-existing brain system (here, the ventral visual stream) from recent evolutionary ancestors, and they either inherited or evolved learning mechanisms that enable individuals to adapt the outputs of that system during their lifespan for word recognition and other core aspects of orthographic processing. According to this view, pre-reading children already possesses many neurons sensitive to letter-like shapes such as T, L, +, etc. that – with supervised learning -- can be simply combined to support invariant word recognition. While we observed only weak single IT neuron tuning for individual letters, we note that such visual encoding is theoretically not the only way that populations of neurons might act as precursors of invariant word recognition behavior, and our IT decoding results empirically demonstrate that here. Regardless of these encoding alternatives, these results suggest that pre-reading children likely have a neural population representation that can readily be re-used to learn invariant word recognition. Relatedly, it has been previously proposed that the initial properties of this system may explain the child’s early competence and errors in letter recognition, e.g. explaining why children tend to make left-right inversion errors by the fact that IT neurons tend to respond invariantly to mirror images of objects (38–40). Over the course of reading acquisition, this neural representation would become progressively shaped to support written word recognition in a specific script. The theory may also explain why all human writing systems throughout the world rely on a universal repertoire of basic shapes (24). As shown in the present work, those visual features are already well encoded in the ventral visual pathway of illiterate primates, and may bias cultural evolution by determining which scripts are more easily recognizable and learnable.

In addition to testing a prediction of this neuronal recycling hypothesis, we also explored the question of how orthographic stimuli are encoded in IT neurons. Decades of research has shown that IT neurons exhibit selectivity for complex visual features with remarkable tolerance to changes in viewing conditions (e.g. position, scale, and pose) (19, 22, 23). More recent work demonstrates that the encoding properties of IT neurons, in both humans and monkeys, is best explained by the distributed complex invariant visual features of hierarchical convolutional neural network models (30, 41, 42). Consistent with this prior work, we here found that the firing rate responses of individual neural sites in macaque IT was modulated by, but did not exhibit strong selectivity to orthographic properties such as letters and letter positions. In other words, we did not observe precise tuning as postulated by “letter detector” neurons, but instead coarse tuning for both letter identity and position. It is possible that, over the course of learning to read, experience-dependent plasticity could fine-tune the representation of IT to reflect the statistics of printed words (e.g. single neuron tuning for individual letters or bigrams). Moreover, such experience could alter the topographic organization to exhibit millimeter-scale spatial clusters that preferentially respond to orthographic stimuli, as have been shown in juvenile animals in the context of symbol and face recognition behaviors (18, 43). Together, such putative representational and topographic changes could induce a reorientation of cortical maps towards letters at the expense of other visual object categories, eventually resulting in the specialization observed in the human visual word form area (VWFA). However, our results demonstrate that, even prior to such putative changes, the initial state of IT in untrained monkeys has the capacity to support many learned orthographic discriminations.

In summary, we found that the neural population representation in IT cortex in untrained macaque monkeys is largely capable, with some supervised instruction, to extract explicit representations of written letters and words. We note that this did not have to be so. Indeed, according to constructivist theories of learning (44), experience determines cortical organization, and thus the visual representations that underlie orthographic processing should be largely determined over developmental time-scales by the experience of learning to read. As such, the IT representation measured in untrained monkeys (or even in illiterate humans) would likely not exhibit the ability to act as a precursor of orthographic processing. Likewise, orthographic processing abilities could have been critically dependent on other brain regions, such as speech and linguistic representations, or putative flexible domain-general learning systems, that evolved well after the evolutionary divergence of humans and Old-World monkeys. Instead, we here report evidence for a “precursor” of visual word form recognition in untrained monkeys. This finding fits with nativist views of cognitive development, according to which learning rests on pre-existing neural representations which it only partially reshapes.

## Methods

### Subjects

The non-human subjects in our experiments were four adult male rhesus macaque monkeys (Macaca mulatta, subjects N, B, S, M). Surgical procedures, behavioral training, and neural data collection are described in detail below. All procedures were performed in compliance with the guideline of National Institutes of Health and the American Physiological Society, and approved by the MIT Committee on Animal Care.

### Visual Images

We randomly subsampled 616 strings (308 words, 308 pseudo-words) from the stimulus set used to test orthographic processing abilities in baboons by Grainger et al. Word strings consisted of four-letter English words, whereas pseudo-word strings consisted of nonsense combinations of four letters, with one vowel and three consonant letters. The entire set of pseudo-words contained bigrams that ranged from those that are very common in the English language (e.g. “TH”) to those that are very uncommon (e.g. “FQ”), as quantified by a broad distribution of English bigram frequency (median = 95, inter-quartile range = 1366; in units of count per million). As such, given the previously established link between bigram frequency and difficulty in lexical decision (17), orthographic stimuli spanned a range of difficulties for the word vs pseudo-word lexical decision task. From these 616 strings, we then generated images of these strings under different variations generative parameters in font size (small/medium/large size) and font case (upper/lower case), fixing the font type (monotype), color (yellow), thus creating a total of 3696 images. We additionally generated images of individual alphabet letters at each of the possibly locations (26 letters x 4 locations x 6 variations in font case/size). We measured IT and V4 responses from passively fixating rhesus macaque monkeys (see below) for a subset of 1120 images from this stimulus set, and used previously measured behavior from trained baboons from the study by Grainger and colleagues (17). Visual images were presented to span 8° of visual angle, with each individual letter of size 0.8°, 1.2, and 1.6° for small, medium, and large variations.

### Baboon behavior

Baboon behavioral data from six guinea baboons performing a lexical decision task was obtained from prior work (17). We focused our analysis on the aforementioned subsampled stimulus set (616 strings).

### Large scale multielectrode recordings

#### Surgical implant of chronic micro-electrode arrays

We surgically implanted each monkey with a head post under aseptic conditions. After behavioral training, we implanted multiple 10 × 10 micro-electrode arrays (Utah arrays; Blackrock Microsystems) in V4 and IT cortex of each monkey. A total of 96 electrodes were connected per array. Each electrode was 1.5 mm long and the distance between adjacent electrodes was 400 µm. Array placements were guided by the sulcus pattern, which was visible during surgery. The electrodes were accessed through a percutaneous connector that allowed simultaneous recording from all 96 electrodes from each array. All behavioral training and testing were performed using standard operant conditioning (fluid reward), head stabilization, and real-time video eye tracking.

#### Eye Tracking

We monitored eye movements using video eye tracking (SR Research EyeLink 1000). Using operant conditioning and water reward, our two subjects were trained to fixate a central white square (0.2°) within a square fixation window that ranged from ±2°. At the start of each behavioral session, monkeys performed an eye-tracking calibration task by making a saccade to a range of spatial targets and maintaining fixation for 500 ms. Calibration was repeated if drift was noticed over the course of the session.

#### Electrophysiological Recording

During each recording session, band-pass filtered (0.1 Hz to 10 kHz) neural activity was recorded continuously at a sampling rate of 20 kHz using Intan Recording Controller (Intan Technologies, LLC). The majority of the data presented here were based on multiunit activity, hence we refer to “neural sites.” We detected the multiunit spikes after the raw data was collected. A multiunit spike event was defined as the threshold crossing when voltage (falling edge) deviated by less than three times the standard deviation of the raw voltage values. In this manner, we collected neural data from macaque V4 and IT from four male adult monkeys (N, B, S, M, weights in kg) in a piecewise manner. We focused our analyses on neural sites that exhibited significant visual drive (determined by p<0.001 comparing baseline activity to visually driven activity); this resulted in 510 IT neural sites and 277 V4 neural sites. Our array placements allowed us to sample neural sites from different parts of IT, along the posterior to anterior axis. However, we did not consider the specific spatial location of the site, and treated each site as a random sample from a pooled IT population. For each neural site, we estimated the repetition-averaged firing rate response in two temporal windows (70ms-170ms and 170ms-270ms after stimulus onset) and concatenated these firing rates for decoding analyses. Single unit analyses focused on the 70ms-170ms time interval.

### Tests of orthographic processing

#### Linear decoders

To test the capacity of ventral stream neural representations to support orthographic processing tasks, we used linear decoders to discriminate between two classes of stimuli (e.g. words versus pseudo-words) using the firing rate responses of neural populations. We used binary logistic regression classifiers with ten-fold cross-validation: decoder weights were learned using the neural population responses to 90% of stimuli and then the performance of the decoder is tested on held-out 10% of stimuli, repeating 10 times to test each stimulus. We repeated this process 10 times with random sampling of neurons. This procedure produces an output class probability for each tested stimulus, and we took the maximum of those as the behavioral “choice” of the decoded neural population.

#### Deep neural network model behavior

We additionally tested a deep neural network model of the primate ventral stream on the exact same images and tasks. We used CORnet-S, a deep recurrent convolutional neural network model that has recently been shown to best match the computations of the primate ventral visual stream (29, 31). CORnet-S approximates the hierarchical structure of the ventral stream, with four areas each mapped to the four retinotopically-defined cortical area in the ventral visual hierarchy (V1, V2, V4, and IT). To do so, we first extracted features from each CORnet-S layer on the same images. As with neural features, we trained back-end binary logistic regression classifiers to determine the ten-fold cross-validated output class probability for each image and for each label.

#### Behavioral metrics

For each behavioral test, we measured the average unbiased performance (or balanced accuracy) as 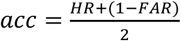, where HR and FAR correspond to the hit-rate and false-alarm-rate across all stimuli. For the lexical decision task, we additionally estimated behavioral patterns across stimuli. To reliably measure behavioral patterns in each individual baboon subject, we grouped the 616 individual stimuli into ten equally-sized bins separately for words and pseudo-words; bins were defined based on the average bigram-frequency of each string in English. We then estimated the average unbiased performance for each stimulus bin using a sensitivity index: *d*’ = *Z*(*HR*) − *Z*(*FAR*) (45), where HR and FAR correspond to the hit-rate and false-alarm-rate across all stimuli within the bin. Across stimulus bins, this resulted in a ten-dimensional pattern of unbiased performances.

#### Behavioral consistency

To quantify the behavioral similarity between baboons and candidate visual systems (both neural and artificial) with respect to the pattern of unbiased performance described above, we used a measure called “*consistency*” 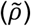 as previously defined (46), computed as a noise-adjusted correlation of behavioral signatures (47). For each system, we randomly split all behavioral trials into two equal halves and estimated the pattern of unbiased performance on each half, resulting in two independent estimates of the system’s behavioral signature. We took the Pearson correlation between these two estimates of the behavioral signature as a measure of the reliability of that behavioral signature given the amount of data collected, i.e. the split-half internal reliability. To estimate the *consistency*, we computed the Pearson correlation over all the independent estimates of the behavioral signature from the model (**m**) and the primate (p), and we then divide that raw Pearson correlation by the geometric mean of the split-half internal reliability of the same behavioral signature measured for each system: 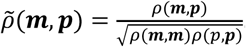. Since all correlations in the numerator and denominator were computed using the same number of trials, we did not need to make use of any prediction formulas (e.g. extrapolation to larger number of trials using Spearman-Brown prediction formula). This procedure was repeated 10 times with different random split-halves of trials. Our rationale for using a reliability-adjusted correlation measure for *consistency* was to account for variance in the behavioral signatures that is not replicable by the experimental condition (image and task).

#### Single neuron analyses

For each neural site, we estimated the selectivity with respect to a number of contrasts (e.g. word vs pseudo-word) using a sensitivity index: 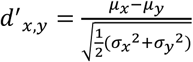 (45). We obtained uncertainty estimates for single neuron selectivity indices by bootstrap resampling over stimuli, and inferred statistical significance using two-tailed exact tests on the bootstrapped distributions. We determined whether neural sites that exhibited significant selectivity for lexical decisions were topographically organized across the cortical tissue using Moran’s *I* (48), a metric of spatial autocorrelation. We compared the empirically measured autocorrelation (averaged over six electrode arrays) to the corresponding distributions expected by chance, obtained by shuffling each electrode’s selectivity 100 times.

